# Neuronal miR-138 represses HSV-2 lytic infection by regulating viral and host genes with mechanistic differences compared to HSV-1

**DOI:** 10.1101/2022.02.28.482433

**Authors:** Siyu Chen, Yue Deng, Hongjia Chen, Yuqi Lin, Xuewei Yang, Boqiang Sun, Dongli Pan

**Affiliations:** State Key Laboratory for Diagnosis and Treatment of Infectious Diseases, The First Affiliated Hospital, Zhejiang University School of Medicine, Hangzhou, Zhejiang, China; Department of Medical Microbiology and Parasitology, Zhejiang University School of Medicine, Hangzhou, Zhejiang, China; innovent Biologics Co., Ltd., Suzhou, Jiangsu 215123, China; Thermo Fisher Scientific, Shanghai 200051, China

**Keywords:** HSV-2, miR-138, ICP0, FOXC1, latency|neurons

## Abstract

Herpes simplex virus 2 (HSV-2) establishes latent infection in dorsal root ganglion (DRG) neurons after productive (lytic) infection in peripheral tissues. A neuron-specific microRNA, miR-138, favors HSV-1 latency by repressing viral *ICP0*, and host *Oct-1* and *Foxc1* genes, yet the role of miR-138 in HSV-2 infection was unknown. The ICP0 mRNAs of HSV-1, HSV-2 and chimpanzee herpesvirus each have one to two canonical miR-138 binding sites. The sites are 100% conserved in 308 HSV-1 and 300 HSV-2 published sequences of clinical isolates. In co-transfection assays, miR-138 repressed HSV-2 ICP0 expression through the seed region and surrounding interactions that are different from HSV-1. An HSV-2 mutant with disrupted miR-138 binding sites on *ICP0* showed increased ICP0 expression in Neuro-2a cells. Photoactivatable ribonucleoside-enhanced crosslinking and immunoprecipitation confirmed miR-138 binding to HSV-2 *ICP0*, and identified *UL19* and *UL20* as additional targets, whose expression was repressed by miR-138 during co-transfection. In Neuro-2a cells, transfected miR-138 and its antagomir decreased and increased HSV-2 replication, respectively, and knockout experiment showed that miR-138’s host targets OCT-1 and FOXC1 were important for HSV-2 replication. In primary mouse DRG neurons, both ICP0 and FOXC1 positively regulated HSV-2 replication, but both overexpressed and endogenous miR-138 suppressed HSV-2 replication primarily by repressing ICP0 expression. Thus, miR-138 can suppress HSV-2 neuronal replication through multiple viral and host pathways. These results reveal functional similarities and mechanistic differences in how miR-138 regulates HSV-1 and HSV-2 infection and indicate an evolutionary advantage of using miR-138 to repress lytic infection in neurons.

**Importance:** Herpes simplex virus 1 (HSV-1) and HSV-2 are closely related viruses with major differences. Both viruses establish latency in neurons from which they reactivate to cause disease. A key aspect of HSV latency is repression of productive infection in neurons. Based on previous work with HSV-1, we investigated the role of a neuron-specific microRNA, miR-138, in HSV-2 infection, and established it as a repressor of HSV-2 productive infection in neuronal cells. This repression is mediated mainly by targeting viral *ICP0* and host *Foxc1* mRNAs, but other pathways also contribute. Despite functional conservation of the role of miR-138 between HSV-1 and HSV-2, many molecular mechanisms differ including how miR-138 represses ICP0 expression and miR-138 targeting of HSV-2 but not HSV-1 *UL19* and *UL20*. To our knowledge, this study provides the first example of host microRNA regulation of HSV-2 infection.

## Introduction

Herpes simplex virus 1 (HSV-1) and HSV-2 are closely related human herpesviruses with important biological and molecular differences. They both establish latent infection in sensory neurons following productive (lytic) infection in peripheral tissues and can reactivate from latency to cause disease (35). HSV-1 mainly infects the facial region and establishes latency in trigeminal ganglia (TG) while HSV-2 usually infects the genitals and establishes latency in sacral dorsal root ganglia (DRG). HSV-1 and HSV-2 can also establish latency in autonomic neurons.

HSV lytic infection proceeds with high expression of lytic genes with immediate-early (IE), early and late genes being expressed in a cascade manner leading to abundant virus replication. Both viral and host factors participate in viral gene expression. Upon entry of the viral genome into the nucleus, viral protein VP16 and host proteins OCT-1 and HCF-1 form a complex that binds to IE promoters and stimulates IE gene expression (17, 26). Meanwhile the IE protein ICP0 counteracts multiple mechanisms of the host intrinsic immunity through its E3 ligase activity (34) thereby stimulating viral replication (10). During latent infection, viral replication stalls and most viral genes are silenced except that active transcription at the latency-associated transcript (LAT) locus drives expression of LATs and some microRNAs (miRNAs) (15, 41, 42, 45). During reactivation, viral gene expression is reverted back to the lytic program in a two-phase process (11, 22). How the virus switches between the lytic and latent programs remains elusive. Epigenetic regulation of transcription appears to contribute significantly to the switch because while heterochromatin is progressively removed during lytic infection (23), it is accumulated during latency establishment (32, 43).

On top of transcriptional regulation, miRNAs regulate gene expression at the post-transcriptional level primarily by binding to 3’ untranslated regions (UTRs) of mRNAs and play important roles in various biological processes including viral infection in mammals (6, 12). HSV encodes nearly 30 miRNAs, a few of which are highly expressed during latency and regulate important viral genes (2-4, 14, 16, 18, 30, 31, 41, 47, 48). Some of them are conserved between HSV-1 and HSV-2. For example, miR-H2 (also known as miR-III for HSV-2) targets ICP0 mRNA (41, 42), and miR-H3 and miR-H4 (also known as miR-I and miR-II, respectively) target the mRNA for the neurovirulence factor ICP34.5 (40, 42). In animal models, one study reported increased neurovirulence and reactivation exhibited by a miR-H2 mutant of HSV-1 in a mouse model (19). However, another study did not recapitulate the results and did not detect a difference in ICP0 expression between the mutant and wild type (WT) (31). HSV-1 miR-H1 and miR-H6 appear to promote lytic infection rather than latency because their deletion impaired reactivation from latency in a mouse model (5). A study using several HSV-2 mutants detected no effect of miR-H2, H3, H4 or H6 on HSV-2 acute replication, latency establishment or reactivation in a guinea pig model (21).

Besides viral miRNAs, some host miRNAs have been shown to influence HSV-1 infection or pathogenesis. For example, miR-155 knockout mice showed increased susceptibility to herpes simplex encephalitis (9). An antagomir of miR-132 reduced angiogenesis following ocular HSV-1 infection (25). miR-101 was reported to repress HSV-1 replication in HeLa cells (49). We previously showed that a neuron-specific miRNA, miR-138, not only represses ICP0 expression (29), but also represses expression of OCT-1 and FOXC1 that are host factors important for HSV-1 lytic infection leading to the hypothesis that miR-138 favors HSV-1 latency through both viral and host pathways (38).

Despite these studies with HSV-1, we have not found any report on regulation of HSV-2 infection by host miRNAs. Interestingly, HSV-2 *ICP0* mRNA was also predicted to be targeted by miR-138. However, the target sites in the *ICP0* 3’ UTR are only partially conserved between HSV-1 and HSV-2, raising doubts about whether miR-138 represses HSV-2 ICP0. Whether the host pathways function during HSV-2 infection was also unknown. To address these questions, as well as the general question of whether host miRNAs participate in regulation of HSV-2 infection, we conducted the following research to investigate the role of miR-138 in HSV-2 infection. The results showed that miR-138 repressed HSV-2 lytic infection in neuronal cells through both viral and host pathways that are largely functionally conserved between HSV-1 and HSV-2. However, multiple mechanistic details in their regulation by miR-138 were different between the two viruses.

## Results

### Predicted miR-138 target sites in HSV-2 *ICP0* 3’ UTR are 100% conserved in sequences of clinical isolates

Although we reported specific high expression of miR-138 in mouse brains and TG, the miR-138 levels in other tissues remained unknown. Therefore, we analyzed miR-138 levels in various mouse tissues and cells and observed substantially higher miR-138 levels in all neuronal cells and tissues including DRG and DRG neurons than in non-neuronal tissues (Fig. 1A), corroborating the notion that miR-138 is a neuron-specific molecule.

**Fig. 1.**
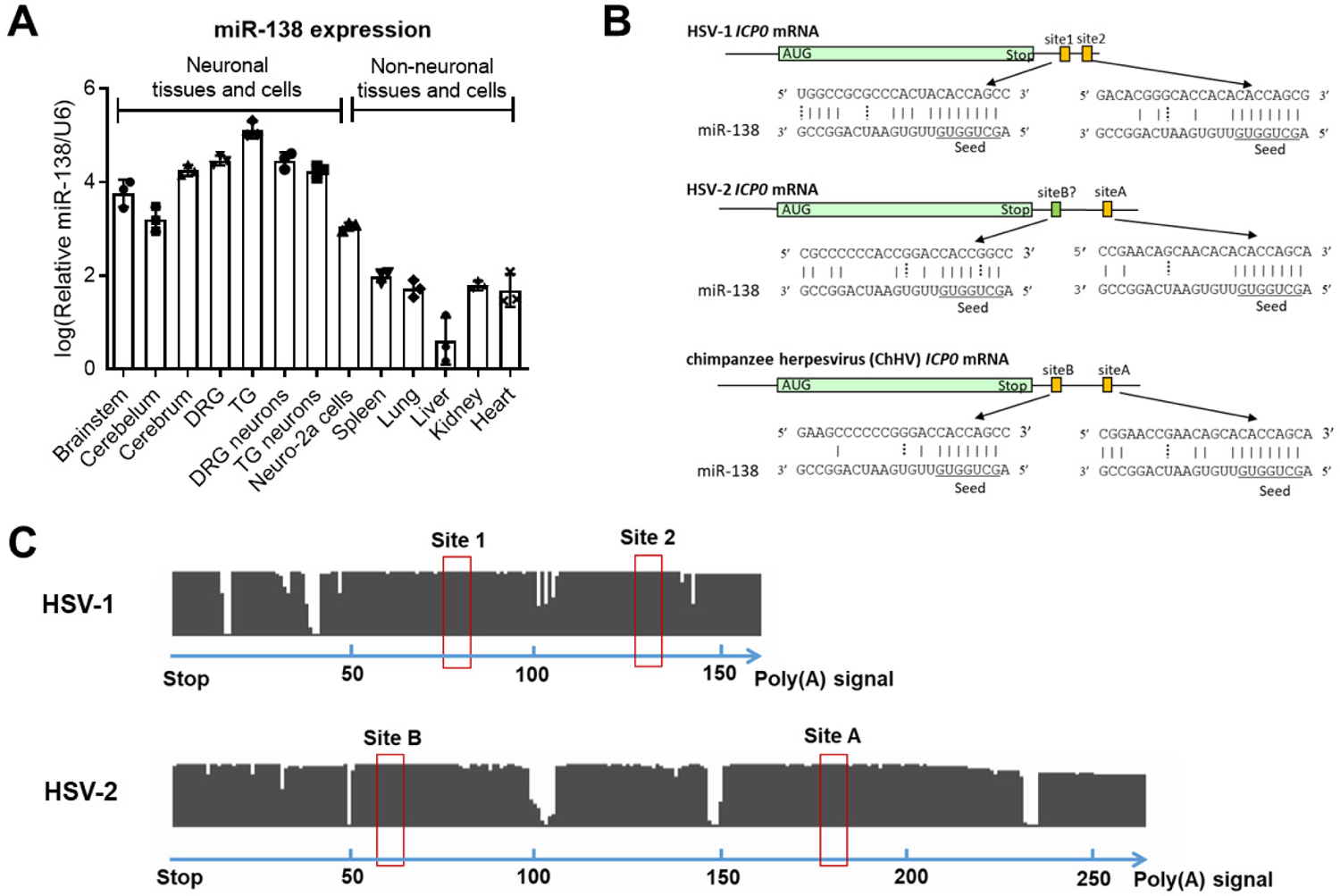
Neuron-specific miR-138 is predicted to have conserved binding sites in HSV-1 and HSV-2 *ICP0* mRNAs. (A) Relative miR-138 levels normalized to U6 levels in the indicated mouse tissues and cells as analyzed by qRT-PCR. n = 3 biologically independent samples. Data are presented as mean values ± standard deviations (S.D.). (B) Graphical representation of predicted binding sites on *ICP0* mRNAs of the three indicated viruses. The miR-138 seed region are underlined. Solid vertical lines represent Watson-Crick base-pairs. Dashed vertical lines represent G-U wobble base pairs. (C) Conservation analysis of the *ICP0* 3’ UTR in HSV-1 (n = 308) and HSV-2 (n = 300) clinical sequences. The clinical sequences were aligned to the sequence from HSV-1 strain 17 (upper) or HSV-2 strain HG52 (lower) by Mega-X. Each gap represents a mismatch, deletion or insertion at a nucleotide position. Putative binding sites of the miR-138 seed region are indicated by red boxes. The numbers at the bottom of each panel represents nucleotide positions.

Sequence inspection reveals two, one and two canonical miR-138 binding sites in the *ICP0* 3’ UTRs of HSV-1, HSV-2 and chimpanzee herpesvirus (ChHV), respectively (Fig. 1B), but no such site in the 3’ UTRs of the *ICP0* homologs of any other alphaherpesviruses we checked including varicella-zoster virus, herpes B virus, saimiriine herpesvirus 1, papiine herpesvirus 2, equid herpesvirus 1, bovine herpesvirus and pseudorabies virus. The partial conservation of the target sites between HSV-1 and HSV-2 is conspicuous in the background of poor conservation elsewhere in the 3’ UTRs (42% sequence identity) that are quite different even in lengths (Fig. 1C). ChHV ICP0 3’ UTR is more similar to HSV-2 (75% sequence identity) than HSV-1. One of the sites in ChHV imperfectly aligns to the HSV-2 canonical site (named siteA). Interestingly, the other site in ChHV imperfectly aligns to a sequence in HSV-2 (named siteB) that nearly perfectly matches the miR-138 seed region except for a mutation that changes an A-U Watson-Crick base pair with miR-138 to a G-U wobble base pair. Therefore we also considered siteB of HSV-2 a putative binding site.

To learn more about the variability of these sites, we checked the National Center for Biotechnology Information (NCBI) database and found 308 HSV-1 and 300 HSV-2 *ICP0* 3’ UTR sequences from clinical isolates and aligned them by the MEGA-X program. Remarkably, we found no mutation in the putative binding sites of the miR-138 seed region in HSV-1 (site 1 and site 2) or HSV-2 (siteA and siteB) despite frequent mutations in the surrounding 3’ UTRs (Fig. 1C), supporting the importance of the miR-138-*ICP0* interaction in HSV biology.

### miR-138 represses HSV-2 ICP0 expression through the 3’ UTR

After preparative work of determining the complete *ICP0* gene sequence of HSV-2 strain 186 (NCBI accession number MT919989) and generation of an antibody against HSV-2 ICP0, we first examined whether miR-138 can regulate HSV-2 ICP0 expression. We constructed an expression plasmid containing the *ICP0* gene with its 3’ UTR (named pV2ICP0) and co-transfected it with a miR-138 or control mimic into 293T cells. Three different control mimics were used: one with a scrambled sequence and two with seed mutations named miR-M138 and miR-M138b (Fig 2A). After co-transfection, western blots revealed repression of ICP0 expression by miR-138 relative to the three controls (Fig. 2B). Interestingly, the extent of the repression, determined to be ∼8-fold by comparisons with dilution series, was the same as the extent to which miR-138 repressed HSV-1 ICP0 expression from a plasmid with the same backbone (Fig. 2C). The effect on HSV-2 ICP0 protein correlated with reduction of *ICP0* mRNA levels by miR-138 which was determined to be ∼5-fold by qRT-PCR (Fig. 2D).

**Fig. 2.**
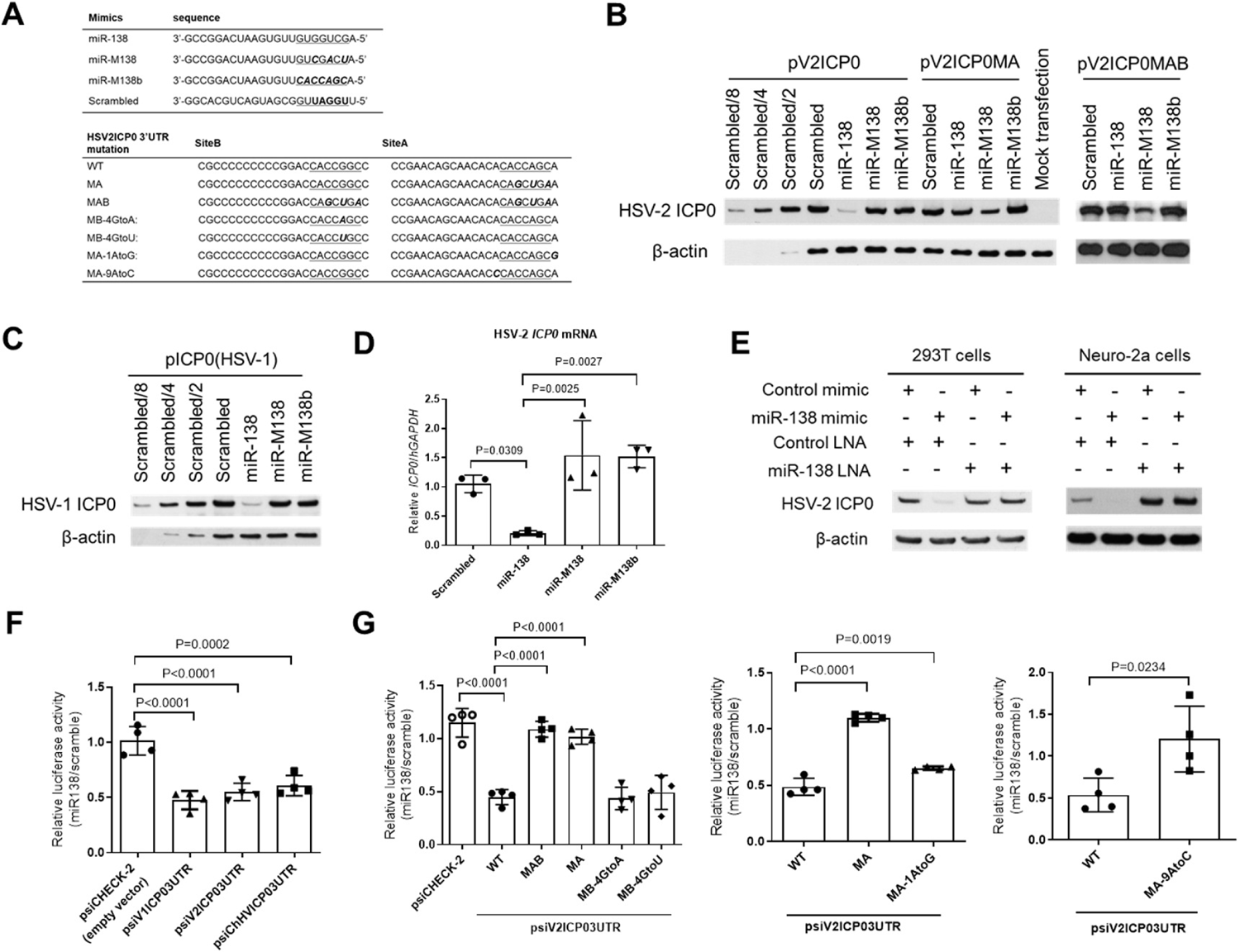
Repression of HSV-2 ICP0 expression by miR-138 and sequence determinants of this repression. (A) Sequences of miRNA mimics and HSV-2 ICP0 3’ UTR mutants. (B-C) 293T cells were co-transfected with 80 ng/ml of the indicated plasmid and 10 nM of the indicated mimic for 30 h before western blot analysis. (D) 293T cells were co-transfected with 20 ng/ml of the indicated plasmid and 10 nM of the indicated mimic for 30 h before qRT-PCR analysis. (E) 293T (left) or Neuro-2a (right) cells were co-transfected with 6 nM of the indicated mimic, 10 nM of the indicated LNA and 60 ng/ml of pV2ICP0 for 30 h (left) or 64 h (right) before western blot analysis. (F-G) 293T cells were co-transfected with 20 ng/ml of the indicated plasmid and 10 nM of a miR-138 or scrambled mimic for 24 h before measurement of luciferase activity. The values are ratios of the activities shown by the miR-138 mimic over those shown by the scrambled mimic. n = 4 biologically independent samples. Data are presented as mean values ± S.D. Data were analyzed by one-way ANOVA with Dunnett’s multiple comparisons tests.

We also co-transfected either a locked nucleic acid (LNA) inhibitor of miR-138 or a negative control LNA together with the mimics. The miR-138, but not control, LNA offset the repressive effect of the miR-138 mimic in 293T cells, confirming that the repression was due to miR-138 (Fig. 2E). To test the effects in neuronal cells, we transfected the mimic-LNA combinations together with pV2ICP0 into Neuro-2a cells and observed repression of HSV-2 ICP0 expression by the miR-138 mimic in the presence of the control but not miR-138 LNA (Fig. 2E). Importantly, in the presence of the control mimic, the miR-138 LNA markedly increased ICP0 expression suggesting that endogenous miR-138 in Neuro-2a cells is sufficient to repress HSV-2 ICP0. As a further test, we constructed dual luciferase constructs with the HSV-1, HSV-2 or ChHV ICP0 3’ UTR downstream of a luciferase gene. Following co-transfection of a miRNA mimic and the luciferase construct into 293T cells, the three 3’ UTRs mediated similar repression of luciferase expression by miR-138 (Fig. 2F). These results demonstrated repression of HSV-2 ICP0 expression by miR-138 through the 3’ UTR.

### Repression of HSV-2 ICP0 requires binding of siteA to the miR-138 seed region and is contributed by interactions outside the seed region

To determine the interactions required for the repression, we first introduced mutations into the pV2ICP0 plasmid. The MA mutations resulted in pV2ICP0MA with three nucleotide substitutions in siteA and the MAB mutations resulted in pV2ICP0MAB with these substitutions in both siteA and siteB (Fig. 2A). The mutations were designed to disrupt binding to the miR-138 seed region but also to complement the mutations in the miR-M138 mimic, such that miR-M138 is complementary to siteA in the ICP0MA mutant and to both siteA and siteB in the ICP0MAB mutant. Still, no sequence match exists between the mutated ICP0 mRNAs and the scrambled or miR-M138b controls. In co-transfection assays in 293T cells, miR-138 no longer repressed ICP0 expression from pV2ICP0MAB and very modestly repressed expression from pV2ICP0MA (Fig. 2B) suggesting that siteA is the major site mediating the repression. As expected, miR-M138 repressed both ICP0MA and ICP0MAB due to the compensatory interactions.

Mutagenesis was then conducted in the luciferase construct to quantitatively assess the sequence determinants of the repression. MA mutations alone removed the repressive effects of miR-138, and MAB mutations had little additional effects (Fig. 2G), in concordance with the importance of siteA but not siteB. Changing the G in siteB that makes the wobble base pair with miR-138 to either A or U to strengthen or disrupt the interaction with miR-138, respectively, had no effect further confirming the dispensability of siteB. However, mutating the A at position t1 (across from position 1 of miRNA) or the A at position t9 of siteA that pairs with the ninth nucleotide of miR-138 both partially alleviated the repression suggesting that interactions surrounding the seed region also contribute to repression. Taken together, repression of HSV-2 ICP0 requires binding of siteA to the miR-138 seed region and is further contributed by interactions surrounding the seed region.

### An HSV-2 mutant with disrupted miR-138 binding sites in *ICP0* showed increased ICP0 expression in Neuro-2a cells

To investigate the role of the miR-138-*ICP0* interaction during HSV-2 infection, we constructed a recombinant virus with the aforementioned MAB mutations. We first replaced nearly the entire *ICP0* gene in strain 186 with *lacZ* by performing homologous recombination and blue-white plaque screening, resulting in an ICP0-null virus designated HSV2ICP0^-^. We then replaced the *lacZ* with either WT *ICP0* or its ICP0MAB mutant, to generate HSV2ICP0R (served as a control) and HSV2ICP0M138 viruses, respectively (Fig. 3A). While HSV2ICP0^-^ showed severe defects in replication, both HSV2ICP0R and HSV2ICP0M138 replicated with WT kinetics in Vero and Neuro-2a cells (Fig. 3B). The lack of a difference in Neuro-2a cells that highly express miR-138 indicated that the impact of the miR-138-ICP0 interaction on HSV-2 replication was limited. Next we investigated ICP0 expression from these viruses. Infection of Neuro-2a cells by these viruses was relatively inefficient so that we could not detect ICP0 protein by western blotting. Therefore we used qRT-PCR to quantify *ICP0* mRNA. Whereas the two viruses showed similar *ICP0* transcript levels in Vero cells, which are known to poorly express miR-138 (29), HSV2ICP0M138 showed significantly higher ICP0 transcript levels than both HSV2ICP0R and WT HSV-2 at 10 hours post infection (hpi) in Neuro-2a cells (Fig. 3C) suggesting that endogenous miR-138 repressed ICP0 expression during HSV-2 infection of Neuro-2a cells even though this did not manifest in observable differences in replication of the viruses.

**Fig. 3.**
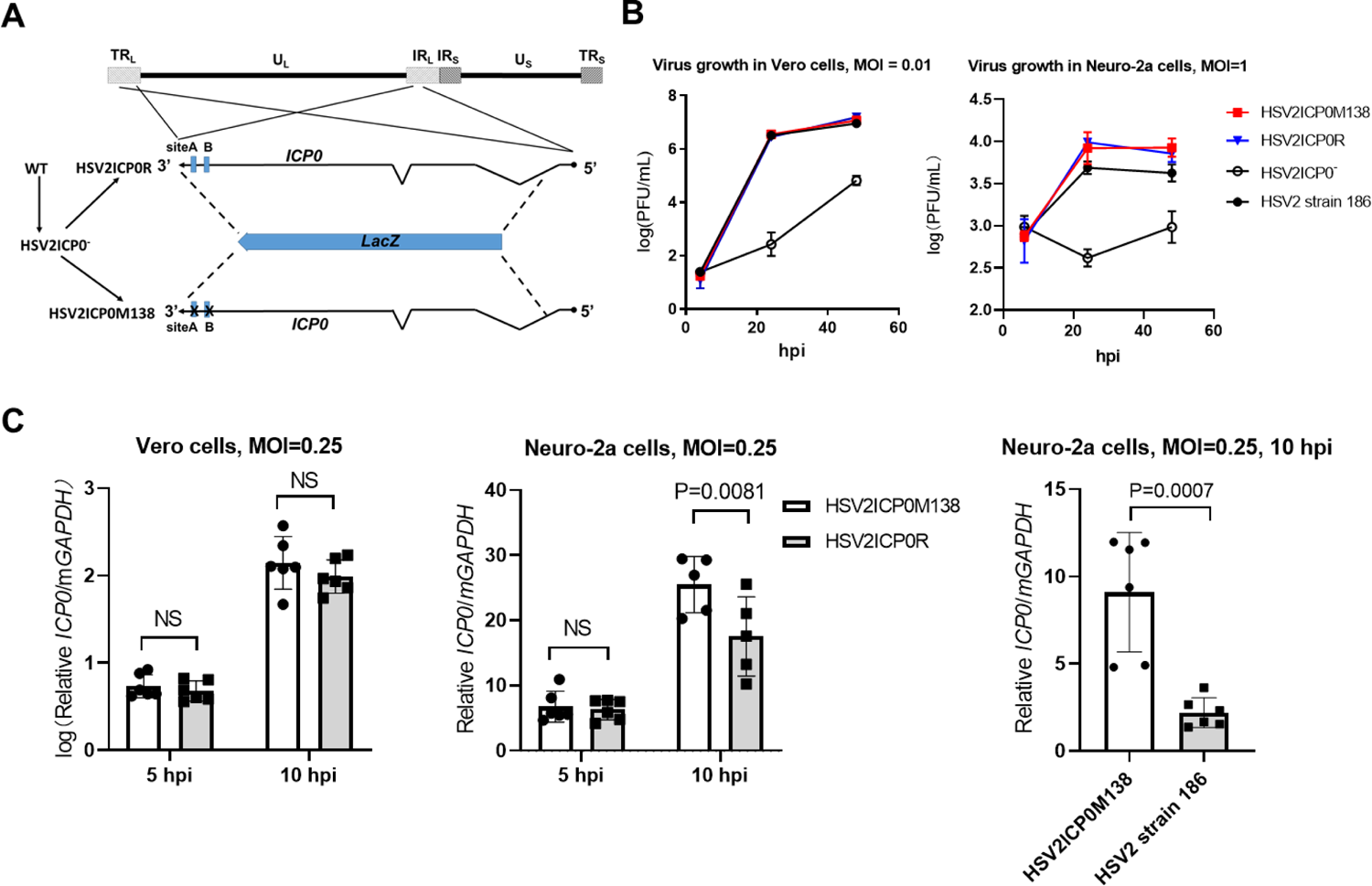
An HSV-2 mutant with altered miR-138 binding sites on ICP0 showed increased ICP0 expression. (A) Schematic diagram of the HSV-2 mutant and rescued virus. The top line represents the HSV-2 genome showing the locations of unique long (UL) and short (US) sequences and repeat long (TRL and IRL) and short (IRS and TRS) sequences. Below is an expanded view of the ICP0 gene present as two copies in two different genomic locations. The zigzag lines represent introns. The blue boxes represent the two putative miR-138 binding sites. The dotted lines represent recombination events that allowed replacement of *ICP0* with *lacZ* (blue arrow), or vice versa. The corresponding virus names are shown to the left with the arrows indicating the order of virus construction. (B) Growth curves of the indicated viruses in Vero (MOI = 0.01) and Neuro-2a cells (MOI = 1). (C) Vero (left) or Neuro-2a (middle and right) cells were infected with the indicated viruses (MOI = 0.25) for the indicated times before qRT-PCR analysis of *ICP0* mRNA levels. n = 3 (B) or 5 or 6 (C) biologically independent samples. Data are presented as mean values ± S.D. Data were analyzed by two-way ANOVA with Sidak’s multiple comparisons tests.

### PAR-CLIP confirmed miR-138 targeting of HSV-2 *ICP0* and identified *UL19* and *UL20* as additional targets

To further confirm the miR-138-*ICP0* interaction and potentially identify additional targets of miR-138 during HSV-2 infection, we performed photoactivatable ribonucleoside-enhanced crosslinking and immunoprecipitation (PAR-CLIP), which detects RNA sequences that cross-link and co-immunoprecipitate with Ago. Previously constructed 293T138 cells that overexpress miR-138 and 293Tcontrol cells (38) were infected with HSV-2 strain 186 for 4 h before PAR-CLIP analysis. The experiment generated sufficient reads for viral target analysis. The experiment was validated by substantial enrichment of PAR-CLIP reads of miR-138 resulting from miR-138 overexpression. Following sequence alignment to the viral genome, 293T138 cells showed 5-fold more reads aligned to ICP0 siteA than 293Tcontrol cells that already showed over 100 reads (Fig. 4A, B), further demonstrating that miR-138 binds to siteA. Although some reads aligned to siteB were detected, they were not enriched by miR-138 overexpression, arguing against binding of miR-138 to siteB.

**Fig. 4.**
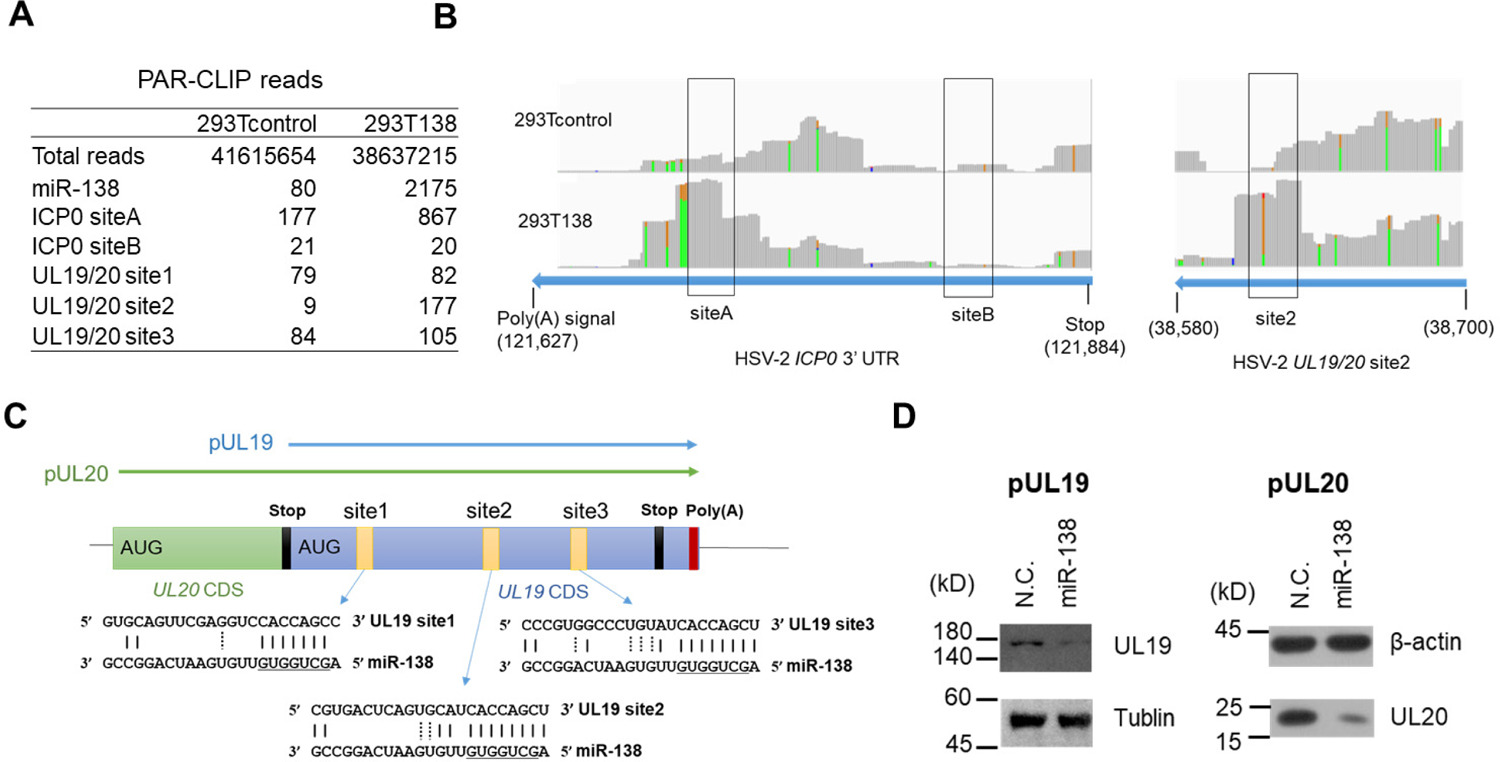
PAR-CLIP identification of HSV-2 targets of miR-138. (A) PAR-CLIP reads aligned to the indicated sites. (B) 293T138 (lower) and 293Tcontrol (upper) cells infected with strain 186 for 4 h were analyzed by PAR-CLIP. The reads coverage peaks were generated in the IGV program. The nucleotide positions are indicated at the bottom. When the fraction of a particular variant is greater than 0.1, the identity of the variant is displayed in color: yellow, A; green G. These A-to-G mutations from 3’ to 5’ ends correspond to T-to-C mutations from 5’ to 3’ ends. (C) Schematic diagram of UL19/20 polycistronic region with green and blue boxes representing the UL19 and UL20 coding sequences (CDS) respectively. The UL19 CDS is also a part of the UL20 3’ UTR. The putative miR-138 binding sites are indicated by small yellow boxes with their sequences shown below. The arrows above the boxes represent the cloned sequences for making pUL19 and pUL20 plasmids. (D) 293T cells were co-transfected with 200 ng/ml of pUL19 (left) or pUL20 (right) plasmid and 20 nM of the indicated mimic for 36 h before western blot analysis using a Flag antibody to detect the Flag-tagged proteins. N.C., negative control.

To examine if there are other viral targets, we checked all the 25 genome locations with mRNA sequences fully complementary to the miR-138 seed region. After applying the criteria of reads > 10 and 293T138/293Tcontrol read ratios > 3, we identified only one location in the *UL19* coding sequence (Fig. 4A, B), which happens to be a part of the *UL20* 3’ UTR too. Interestingly, this region contains three canonical sites for miR-138 (Fig. 4C) although only one (site 2) was detected by PAR-CLIP. We constructed a UL19 expressing plasmid containing its whole coding region as well as a UL20 expressing plasmid containing its 3’ UTR (Fig. 4C). Co-transfection assays showed repression of both UL19 and UL20 by miR-138 (Fig. 4D) suggesting that miR-138 also targets these viral mRNAs in addition to *ICP0*.

### ICP0-dependent and independent suppression of HSV-2 replication and gene expression by miR-138

To examine the overall impact of miR-138 on HSV-2 replication, we transfected the miRNA mimics into Neuro-2a cells before HSV-2 infection. The mutant viruses described above were utilized to assess whether the effects were dependent on ICP0. Relative to two negative control mimics, the miR-138 mimic significantly suppressed replication of both HSV2ICP0M138 and HSV2ICP0R (Fig. 5A), suggesting that the effects were at least partially independent of the miR-138-*ICP0* interaction. On the other hand, the effects on HSV2ICP0R (∼6-fold) were greater than those on HSV2ICP0M138 (∼3.4-fold), and HSV2ICP0M138 yields were significantly higher than HSV2ICP0R yields in miR-138 transfected cells indicative of a contribution from the miR-138-*ICP0* interaction too. The ICP0-independent suppression was also demonstrated by the observation that transfected miR-138 also reduced HSV2ICP0^-^ yields (Fig. 5B). To determine the impact of endogenous miR-138, we transfected a miR-138 antagomir into Neuro-2a cells before infection. Relative to the control antagomir, the miR-138 antagomir modestly (∼2-fold) but significantly increased the yields of HSV2ICP0M138 and HSV2ICP0R, as well as WT HSV-2, suggesting that endogenous miR-138 also suppressed HSV-2 replication independently of ICP0. To investigate the effects of miR-138 on viral gene expression, we analyzed *ICP0* (immediate-early), *TK* (early) and *UL19/20* (late) mRNA levels at 5 and 18 hpi after transfection with miRNA mimics. Compared to the control mimic, the miR-138 mimic had no significant effect at 5 hpi but significantly reduced levels of all the three transcripts at 18 hpi. Although direct regulation by miR-138 could contribute to the reduction of *ICP0* and *UL19/UL20* levels, the effects on *TK* which is not targeted by miR-138 and should not be regulated directly by UL19/UL20 indicate secondary effects of regulating ICP0 or other targets of miR-138. We also tested the role of miR-138 in some non-neuronal cell types. Transfected miR-138 suppressed HSV-2 replication independently of ICP0 in HeLa cells but had no observable effect in 293T or Vero cells (Fig. S1) suggesting that the impact of miR-138 on HSV-2 infection is cell-type or host-species specific.

**Fig. 5.**
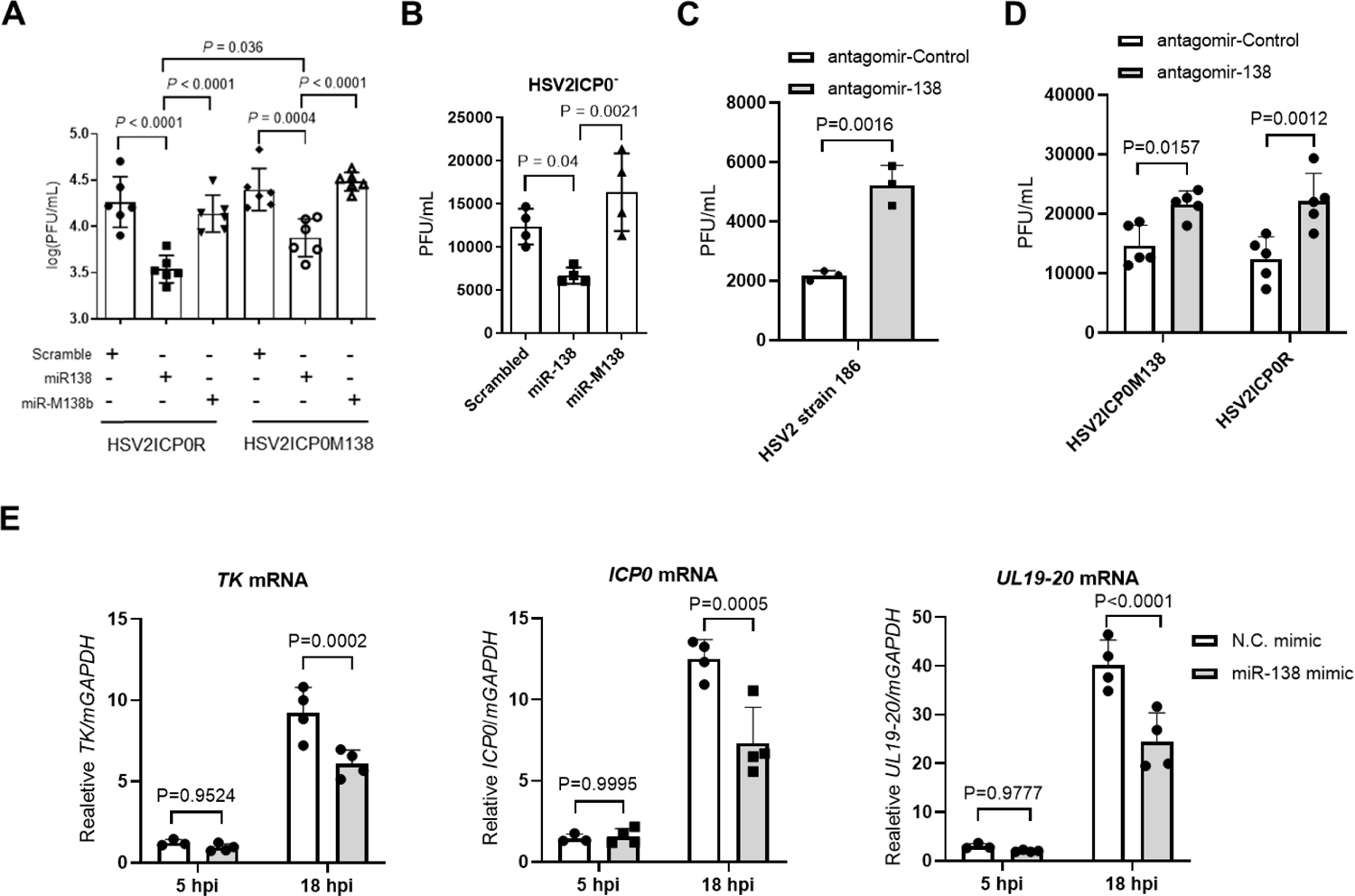
ICP0-dependent and independent suppression of HSV-2 replication and gene expression by miR-138 in Neuro-2a cells. (A) Neuro-2a cells were transfected with 20 nM of the indicated mimic for 24 h, then infected with the indicated virus for 48 h (MOI = 0.2) before viral titer measurements. n=6 (B) Same as A, but HSV2ICP0^-^ virus was used for infection (MOI = 5). n=4 (C) Neuro-2a cells were transfected with 160 nM of the indicated antagomir for 24 h, then infected with HSV-2 strain 186 for 24 h (MOI = 0.4) before viral titer measurements. n = 3 (D) Same as C, but the indicated viruses were used for infection (MOI = 1). n = 5 (E) Neuro-2a cells were infected with the HSV-2 strain 186 (MOI = 0.2) for the indicated times before qRT-PCR analysis of *ICP0, TK* and *UL19-20* mRNA levels. n = 4. Data are presented as mean values ± S.D. Data were analyzed by one-way ANOVA with Dunnett’s multiple comparisons tests (A, B), two-tailed unpaired *t*-tests (C) or two-way ANOVA with Sidak’s multiple comparisons tests (D,E).

### Host targets of miR-138, OCT-1 and FOXC1, are important for HSV-2 replication in Neuro-2a cells

Since the viral targets could not account for the full impact of miR-138 on HSV-2 replication, we investigated host targets. Our previous study showed that FOXC1 and OCT-1 are host targets of miR-138 that can mediate suppression of HSV-1 replication by miR-138 (38). We confirmed that during HSV-2 infection of Neuro-2a cells, transfected miR-138 reduced levels of *Foxc1* and *Oct-1* mRNAs (Fig. S2). Quantification of copy numbers in this experiment showed that endogenous *Oct-1* mRNA is more abundant than *Foxc1* mRNA. To examine the roles of these host transcription factors in HSV-2 infection, we used FOXC1 and OCT-1 knockout cells that were constructed previously (38) and in this study, respectively. The yields of WT strain 186 in all these knockout cells were lower than in the parental Neuro-2a cells in concordance with the supportive role of both OCT-1 and FOXC1 in HSV-2 replication (Fig. 6A, B). While transfected miR-138 still significantly repressed HSV-2 replication in OCT-1 knockout cells, it no longer did so in two independently constructed FOXC1 knockout cell lines further substantiating the importance of FOXC1 in mediating miR-138 suppression of HSV-2 replication. Moreover, transfected human FOXC1 increased HSV-2 replication in Neuro-2a cells and rescued the impairment in replication in FOXC1 knockout cells (Fig. 6C). Therefore, both FOXC1 and OCT-1 are required for efficient HSV-2 replication in Neuro-2a cells with a possible more important role from FOXC1 in mediating suppression by miR-138.

**Fig. 6.**
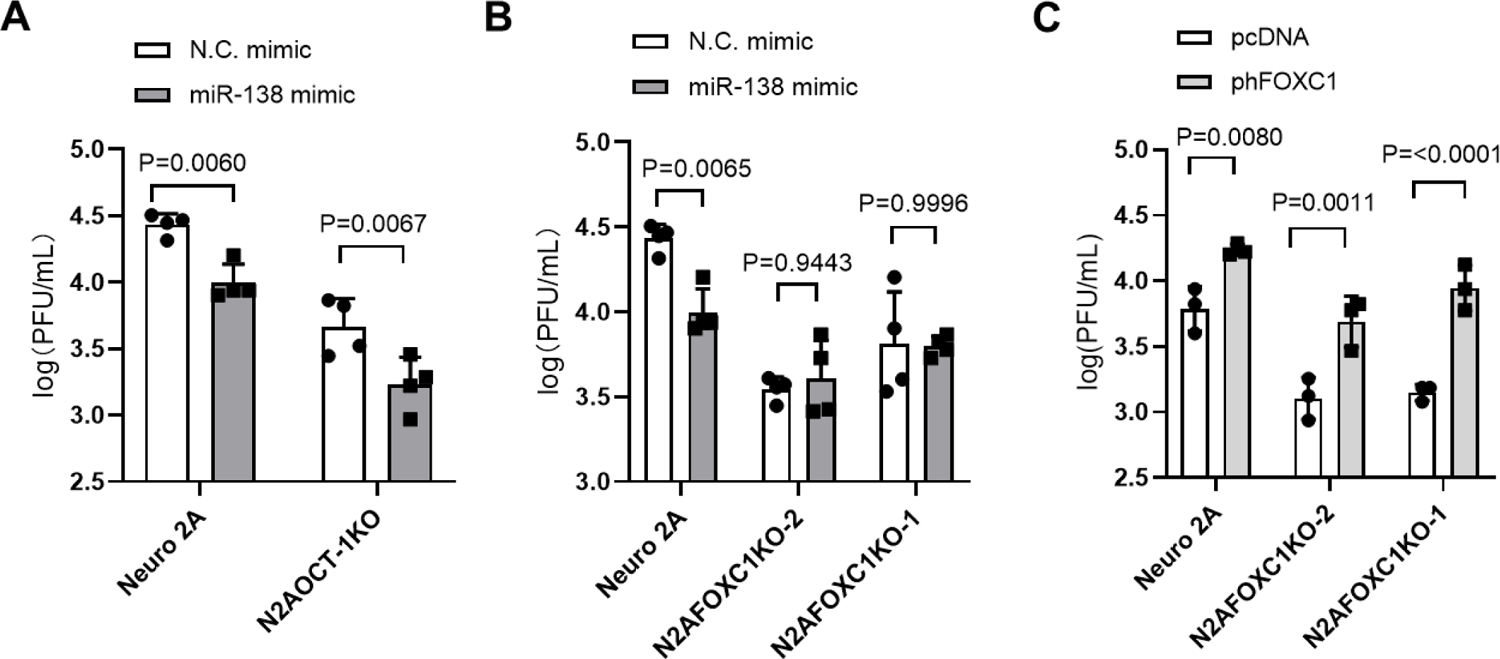
FOXC1 and OCT-1 are important for HSV-2 replication in Neuro-2a cells. (A, B) The indicated cell lines were transfected with 10 nM of the indicated mimic for 24 h and infected with WT 186 virus for 48 h (MOI = 0.2) before viral titer measurements. (C) The indicated cell lines were transfected with 200 ng/ml of the indicated plasmid for 24 h and infected with WT 186 virus for 48 h (MOI = 0.2) before viral titer measurements. Data are presented as mean values ± S.D. n = 4 (A, B) or 3 (C) biologically independent samples. Data were analyzed by two-way ANOVA with Sidak’s multiple comparisons tests.

### ICP0-dependent suppression of HSV-2 replication by miR-138 and the role of FOXC1 in mouse DRG neurons

Since DRG are major tissues of HSV-2 latency, we cultured primary mouse DRG neurons and used them to examine the effects of ICP0, human FOXC1 and miR-138. HSV-2 replication in these neurons was inefficient and often undetectable at low MOIs. At an MOI of 5, HSV2ICP0^-^ exhibited a modest but significant replication defect (Fig. 7A) consistent with the importance of ICP0.

**Fig. 7.**
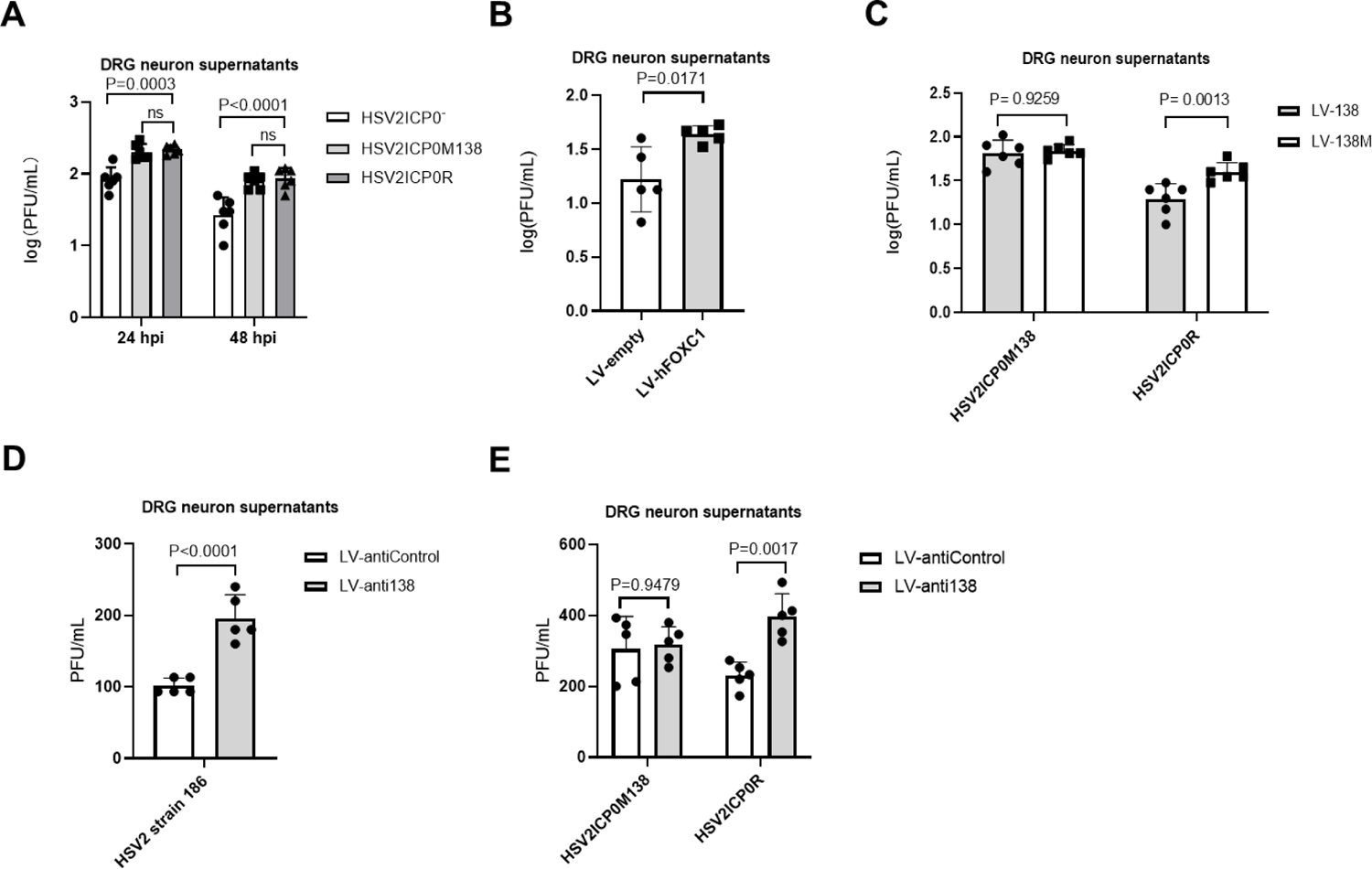
Effects of ICP0, FOXC1 and miR-138 on HSV-2 replication in mouse DRG neurons. (A) DRG neurons were infected with the indicated viruses (MOI = 5). Supernatants were collected at the indicated times and titrated. (B) DRG neurons were transduced with lentivirus expressing FOXC1 or the control lentivirus for 4 days before infection with WT strain 186 (MOI = 5). Supernatants were collected at 24 hpi and titrated. (C) DRG neurons were transduced with lentivirus expressing miR-138 (LV-138) or a control lentivirus (LV-138M) for 4 days and then infected with HSV-2 strain 186 (MOI = 5). Supernatants were collected at 24 hpi and titrated. (D) DRG neurons were transduced with a lentivirus that expresses antisense sequences antagonizing miR-138 (LV-anti138) or a control lentivirus (LV-antiControl) for 4 days and then infected with HSV2 strain 186 (MOI = 5). Supernatants were collected at 24 hpi and titrated. (E) Same as D, but the indicated viruses were used for infection (MOI = 5) Supernatants were collected at 24 hpi and titrated. Data are presented as mean values ± S.D. n = 6 (A, B) or 5 (C, D and E) biologically independent samples. Data were analyzed by two-way ANOVA with Sidak’s multiple comparisons tests (A, C and E) or two-tailed unpaired *t*-tests (B,D).

The supportive role of FOXC1 was also evident because transduction of these neurons with lentivirus expressing FOXC1 caused increased HSV-2 yields compared to control lentivirus (Fig. 7B). To examine the role of the miR-138-*ICP0* interaction in DRG neurons, we compared HSV2ICP0M138 and HSV2ICP0R. Although they showed no difference in replication in untreated DRG neurons (Fig. 7A), transduction of the neurons with lentivirus expressing miR-138 significantly reduced replication of HSV2ICP0R but not HSV2ICP0M138 virus (Fig. 7C). Conversely, transduction of the neurons with lentivirus expressing antisense sequences targeting miR-138 significantly increased replication of WT HSV2 and HSV2ICP0R but not HSV2ICP0M138 virus (Figs. 7D and 7E). Thus, while both ICP0 and FOXC1 can promote HSV-2 replication meaning that targeting them should contribute to suppression of HSV-2 replication, targeting ICP0 appears to be more important for this suppression in mouse DRG neurons.

## Discussion

Based on our related study with HSV-1, we extended our analyses and showed repression of HSV-2 lytic infection by miR-138 in neuronal cells. A major mechanism of this repression is the interaction between miR-138 and *ICP0* mRNA as demonstrated by repression of ICP0 expression by miR-138 in transfected and infected cells as well as the necessity of this interaction for miR-138 suppression of HSV-2 replication in DRG neurons. The importance of this interaction is also reflected by its conservation within closely related human and chimpanzee viruses, especially when this conservation occurs in the background of poor conservation between HSV-1 and HSV-2 3’ UTR sequences outside the binding sites of the miR-138 seed region. Interestingly, the ICP0 3’ UTRs are more conserved between HSV-2 and ChHV, consistent with higher overall sequence identity between HSV-2 and ChHV than between HSV-1 and HSV-2 (37), which has led to a hypothesis where HSV-1 resulted from ancient divergence whereas HSV-2 arose from cross-species transmission from ancestors of modern chimpanzees (46). One major difference between HSV-1 and HSV-2 is that siteA in HSV-2 has an A at position t1 that is absent in both HSV-1 *ICP0* sites. Such an A is known to enhance the efficacy of repression due to its binding to a pocket in Ago protein (36). Indeed we show that this A contributes significantly to repression, which may help explain why miR-138 represses HSV-2 ICP0 as strongly as it represses HSV-1 ICP0 despite one fewer canonical site in HSV-2 than HSV-1 *ICP0*. On the other hand, contribution from base pairing at position t9 is unexpected since this position is usually thought to be dispensable (6). However, it may not be coincidental that four out of five canonical sites in HSV-1, HSV-2 and ChHV *ICP0* 3’ UTRs have this base pairing at position t9, raising the possibility that base pairing beyond the seed region may sometimes contribute to gene repression.

Besides ICP0, multiple other viral and host targets of miR-138 positively regulate HSV-2 replication. An interaction not conserved with HSV-1 is miR-138 targeting of HSV-2 *UL19* and *UL20*. *UL19* encodes the essential capsid protein VP5 (13), and *UL20* is a membrane protein involved in cytoplasmic virion envelopment and infectious virus production (1). Therefore these viral targets are also important for the lytic cycle. An important host target of miR-138 is *Foxc1*, which encodes a transcription factor that stimulates HSV-1 gene expression in neuronal cells by antagonizing host epigenetic silencing mechanisms (38). Similar to its role in HSV-1 infection, FOXC1 can enhance HSV-2 replication in both Neuro-2a cells and mouse DRG neurons, and is critical for miR-138 suppression of HSV-2 replication in Neuro-2a cells. We also show that another host target of miR-138, OCT-1, is important for HSV-2 replication in Neuro-2a cells. Taken together, there are multiple pathways through which miR-138 represses HSV-2 neuronal replication.

We have already tested HSV2ICP0M138 and HSV2ICP0R viruses in a mouse model of corneal inoculation (data not shown). However, the model was unsatisfactory in that the results showed weak gene expression during acute infection and no evidence of establishment of latency in TG, so the data could not be interpreted. Future studies need to examine a different model such as a mouse or guinea pig model of genital inoculation which better resembles natural infection and has been successfully used for studying HSV-2 latency and reactivation (7, 21, 39). While its role in HSV-2 latency has yet to be elucidated in vivo, miR-138 suppression of HSV-2 replication especially in DRG neurons where HSV-2 latency occurs as well as its repression of multiple genes important for the lytic cycle is consistent with a conducive role of miR-138 in HSV-2 latency similar to its role in HSV-1 latency.

Under selection pressure, viruses have evolved to adapt to different host species and cells. The similarities and differences in how miR-138 regulates HSV-1 and HSV-2 infection may reflect adaption of the two viruses to similar yet not exactly the same neuronal environment as they have been reported to prefer different neuron subsets for latency (8, 24). It is notable that the interaction between miR-138 and *ICP0* mRNA or its homologs is conserved only in a few human and ape alphaherpesviruses. Since alphaherpesviruses have generally evolved to establish latency in neurons, it will be worth investigating whether other neuronal miRNAs are exploited by these viruses to favor latency.

## Materials and Methods

### Cells

Vero, Neuro-2a, U2OS, HeLa and 293T cells were obtained from American Type Culture Collection. These cells were maintained in Dulbecco’s Modified Eagle Medium supplemented with 5% fetal bovine serum and 1% penicillin– streptomycin (Gibico) in 5% CO_2_ at 37 °C. Mice were housed in accordance to the guidelines for the Care and Use of Medical Laboratory Animals (Ministry of Health, China) and approved by the Animal Research Committee of Zhejiang University. Dorsal root ganglion (DRG) neurons were isolated and cultured as described previously with slight modifications (20). Briefly, 6-week-old CD-1 (Institute for Cancer Research) male mice were anaesthetized with isoflurane (RWD) for 1 min and transcardially perfused with approximately 10 ml of phosphate buffered saline (PBS). DRG were dissected and digested in collagenase/dispaseII solution (Sigma-Aldrich) at 37 °C for 1 h. Neurons were purified by gradient separation in an OptiPrep gradient (Sigma-Aldrich) followed by two washes with Neurobasal-A medium (Thermo Fisher Scientific) + 2% B27 (Gibco). Purified neurons were counted and plated on 10-mm coverslips pretreated with poly-D-lysine (Sangon Biotech) and laminin (Sigma-Aldrich). About 6,000 neurons were plated on each coverslip. After the neurons adhered, the coverslips were transferred to a 48-well plate and cultured in Neurobasal-A medium + 2% B27 + 50 ng/ml of neurturin (R&D Systems) + 50 ng/ml of neuronal growth factor (R&D Systems) + 50 ng/ml of glial-derived neurotrophic factor (R&D Systems) + 1 μg/ml of mitomycin C (MedChemExpress) for 3 to 4 days before being used.

### Plasmids

To construct pV2ICP0WT, the HSV-2 ICP0 gene and flanking sequences were amplified using V2ICP0Ecof and V2ICP0Hindr1 primers and the WT strain 186 genome as the template, and inserted between EcoRI and HindIII sites of pUC18 (Addgene). M138A and M138AB mutations were introduced using the overlapping PCR method. Briefly, using pV2ICP0WT as a template, fragment ab was amplified using V2ICP0Pstf and V2-0M138Af primers, and fragment cd was amplified using V2-0M138Ar and V2ICP0Hindr1 primers. The overlapping ab and cd fragments were mixed in a 1:1 molar ratio to serve as templates and V2ICP0Pstf and V2ICP0Hindr1 served as primers to amplify a fragment that was inserted between PstI and HindIII sites of pV2ICP0WT resulting in pV2ICP0M138A. pV2ICP0M138AB was constructed in the same way on the basis of pV2ICP0M138A using pV2-M138Bf and pV2M138Br as primers. pV2ICP0long and pV2ICP0longMAB were constructed in the same way except that the initial cloning step used V2ICP0Ecof and V2ICP0Hindr2 primers to amplify the ICP0 gene with longer flanking sequences that were inserted into pUC18. pICP0(HSV-1) was the same as the pRS-1 plasmid previously described (29). FLAG-HA-pcDNA3.1^-^ was obtained from Addgene and renamed pcDNA in this study. To construct pUL19 and pUL20, the UL19 and UL20 gene sequences were amplified using pUL19-F/R and pUL20-F/R primers, respectively, and inserted between EcorRI and HindIII sites of pcDNA. FOXC1 expressing plasmid pFOXC1human, here named phFOXC1, also has the pcDNA backbone and was previously described (38). To construct the psiV1ICP03UTR luciferase construct, the HSV-1 ICP0 3’UTR was amplified from pICP0(HSV-1) using psiV1ICP0UTRf and psiV1ICP0UTRr primers and inserted into psiCHECK-2 (Promega). psiChHVICP03UTR was constructed by inserting the synthesized ChHV ICP0 3’ UTR into the same position. To construct psiV2ICP03UTRWT, psiV2ICP03UTRMA and psiV2ICP03UTRMAB, the HSV-2 ICP0 3’UTR was amplified from WT pV2ICP0, pV2ICP0MA and pV2ICP0MAB, respectively, using psiV2ICP0UTRf and psiV2ICP0UTRr primers and inserted into psiCHECK-2. For luciferase constructs with other mutations, mutations were introduced by the Q5 Site-Directed Mutagenesis kit (New England Biolabs) according to the manufacturer’s protocol. For lentivirus expression of FOXC1, the FOXC1 gene was amplified from the phFOXC1 plasmid using LVFoxc1Fw and LVFoxc1Rv primers and inserted between the EcoRI and BamHI sites of the pLVX-EF1α-IRES-mCherry (renamed pLV-empty here, obtained from Biofeng) resulting in pLVFOXC1. All the primer sequences are listed in Table S1.

### Viruses

HSV-2 strain 186 was kindly provided by Donald M. Coen (Harvard Medical School). Recombinant viruses were constructed by homologous recombination in transfected cells as previously described (27). Briefly, to construct HSV2ICP0^-^, the *lacZ* gene was PCR amplified from the TKLTRZ virus (44) genome using lacZfwAfl and lacZrvSma primers and inserted between the AflII and PsiI restriction sites of the pV2ICP0long plasmid to generate pV2ICP0lacZ. pV2ICP0lacZ was linearized with ScaI and transfected together with the purified genome of strain 186 into U2OS cells. When cytopathic effects were observed, the supernatant was subjected to plaque purification and X-gal staining. Single blue plaques were picked and expanded before another round of plaque purification. After three rounds of plaque purification the viruses were screened further by PCR and sequencing to select for single plaques with both *ICP0* copies replaced with *LacZ*. These viruses were then propagated. Lack of ICP0 expression was further verified by western blots after infection of Vero cells. One of the verified viruses were named HSV2ICP0^-^ and used for this study. To construct HSV2ICP0M138 and HSV2ICP0R viruses, the viral genome was purified from HSV2ICP0^-^ virus and transfected together with linearized pV2ICP0longMAB or pV2ICP0long plasmids into Vero cells, respectively. The resulting viruses were plaque purified and screened by PCR, sequencing and western blotting as described above.

### Quantitative RT-PCR

To quantify miR-138 levels in cells or tissues, total RNA was purified using an Eastep super total RNA extraction kit following the protocol for retaining small RNAs provided by the manufacturer. Reverse transcription using stem-loop primers and subsequent PCR were carried out using the TaqMan MicroRNA Assay kit, TaqMan MicroRNA RT kit and TaqMan Universal PCR master Mix II no UNG (all from Applied Biosystems). miRNA levels were quantified by using standard curves generated from serially diluted synthetic miRNAs. miR-138 levels were normalized to U6 transcript levels. To quantify HSV-2 *ICP0* transcript, RNA were isolated using an Easy RNA Extraction kit (Easy-Do Biotech), reverse transcription was conducted using a HiScript II Q Select RT SuperMix kit and PCR was performed using a ChamQ Universal SYBR qPCR kit (all from Vazyme Biotech). *ICP0* transcript levels were normalized to mouse GAPDH transcript levels. All primer sequences are listed in Table S2.

### Sequencing and sequence alignment

The complete ICP0 gene sequence of HSV-2 strain 186 was determined by Sanga sequencing of pV2ICP0WT using the primers listed in Table S3. Alignment of clinical sequences was conducted using the Molecular Evolutionary Genetics Analysis (MEGA-X) multiple sequence alignment program. Clinical isolate sequences were obtained from NCBI databases (see Table S4). The reference sequence of HSV-1 was from strain 17 (GenBank accession number NC_001806), and that of HSV-2 was from strain HG52 (GenBank accession number NC_001798.2).

### Transfection

Plasmids and RNA oligonucleotides were transfected into cells in 24-well plates using Lipofectamine 3000 (Thermo Fisher) according to the manufacturer’s instructions. Synthetic miRNA mimics (miRCURY LNA miRNA mimics) and LNAs (miRCURY LNA miRNA power inhibitors) were obtained from QIAGEN. Synthetic antagomir were obtained from Ribobio. The final concentrations used during transfection were described in figure legends.

### Western blot analysis

Western blotting was performed as described previously (28). Rabbit polyclonal antibody for HSV-2 ICP0 was raised against the synthetic peptide SRADPGPERPPRQTP-Cys corresponding to amino acid residues 9 to 23 of the protein by Sangon Biotech. The antibody was used at a dilution factor of 2000. The following other primary antibodies and dilutions were used: anti-β-tubulin antibody (1:5,000, Tianjin Sungene Biotech), mouse anti-FLAG antibody (1:1,000, Sigma-Aldrich); anti-β-actin antibody (1:10,000, Sigma-Aldrich). HRP-conjugated goat anti-mouse, goat anti-rabbit and rabbit anti-goat antibodies (SouthernBiotech) were used as secondary antibodies with a dilution of 1:2,000.

### Dual luciferase assays

Luciferase activities were detected by dual-luciferase assay kit (Yeasen Biotech). The Renilla luciferase activity was normalized to the firefly luciferase activity and then the miR-138 group normalized to the scramble group.

### PAR-CLIP

PAR-CLIP was performed as described using previously generated 293T138 and 293Tcontrol cells (38), except that after addition of 4-thiouridine, the cells were infected with WT HSV-2 strain 186 for 4 h at an MOI of 1 before crosslinking. Sequencing was performed using a HiSeq 2500 sequencer (Illumina) by single-end sequencing of 50 cycles. Reads were trimmed to discard adapters, low-quality reads and reads less than 20 bases long using fastx_clipper version 0.0.13 (http://hannonlab.cshl.edu/fastx_toolkit/). Reads containing the sequence of the binding site of the miR-138 seed region (CACCAGC) were obtained using the “grep” command. The HG52 genome (GenBank accession number NC_001798.2) was used because the complete genome sequence of strain 186 is not available. To avoid confounding of sequence alignment by the repeat regions, a HSV-2 genome fasta file, named HSV2norepeat was created by removing the terminal repeat regions (TRL and TRS) from the HSV-2 strain HG52 genome. The corresponding transcript annotation file was created manually according to the gff format based on the coordination information from GenBank. The human genomic files as well as the corresponding transcript annotation files (in the gtf format) were downloaded from the Harvard Medical School Research Computing server (https://rc.hms.harvard.edu). PAR-CLIP reads were aligned to the human or HSV-2 genome using Tophat2 version 2.1.0 (https://ccb.jhu.edu/software/tophat/index.shtml). Read counts for each transcript were determined by Htseq-count version 0.9.1 (https://htseq.readthedocs.io/). The putative target sites within the viral genome were counted manually after the alignment files (in bam format) were loaded into IGV version 2.3.91 (tp://software.broadinstitute.org/software/igv/). The coverage plots for reads aligned to the ICP0 3’ UTR and to the region surrounding UL19/UL20 site2 were also generated in IGV.

### Construction of N2AFOCXKO and N2AOCT1KO cells

N2AFOXC1KO cell lines were described previously (38). To knock out OCT-1, we targeted a sequence of OCT-1, TTTCTGAAAGTCCAGACCAT, corresponding to nucleotides 107-126 of the mouse Foxc1 coding region. Synthetic oligonucleotides were designed as described (33) and cloned into the PX459 vector (Addgene). Two hundred ng of the resulting plasmid was transfected into 1.0 × 10^5^ Neuro-2a cells in a 24-well plate. After 24 h, the supernatant was replaced with fresh medium and cells were transfected again. At 24 h after the second transfection, the medium was replaced with fresh medium containing 1 μg/mL of puromycin. Forty-eight h later, cells were washed with PBS and trypsinized. One portion of the cells was used for a T7 endonuclease 1 assay (33), which showed successful editing at the desired location (data not shown). The other portion was diluted and seeded in 96-well plates with a density of 0.5 cells per well in the presence of 0.5 μg/ml of puromycin. Wells with single cells were labelled after 12 h of complete adherence. When cells were confluent, they were trypsinized and transferred to a 24-well plate for expansion. When the cells were confluent again, a portion was used for further expansion in medium with 0.5 μg/ml of puromycin. The remaining cells were used for genomic DNA extraction using the Tissue & Cell Genomic DNA Purification Kit (GeneMark), followed by PCR using the primers ATCCTCCCTCCTACAGC and AGCATTCTCAGTTATGGC and then sequencing using these primers, one colony gave clean sequencing data, with a deletion of 7 nucleotides. The expanded cell line was designated N2AOCT1KO.

### Lentivirus transduction of DRG neurons

The lentiviruses expressing miR-138 (LV-138) and its antisense sequences (LV-anti138) as well as their respective controls (LV-138M and LV-antiControl) were previously described (38). The lentivirus expressing human FOXC1 (LV-hFOXC1) and its control lentivirus (LV-empty) were produced as previously described using pLV-hFOXC1 and pLV-empty, respectively. Lentivirus titers were determined by qRT-PCR amplification of a region in the mCherry gene within the lentivirus genome. DRG neurons were isolated and seeded on 10-mm cover slips in a 48-well plate for 3 days before being transduced with 6 x 10^7^ genome copies of lentivirus per well. Four days after transduction, the neurons were infected with HSV-2.

### Statistical analyses

Statistical analyses were performed using GraphPad Prism version 8.0.2 (GraphPad Software). T tests were used for comparing two groups. For multiple comparisons, one-way ANOVA with Dunnett’s multiple comparisons tests or two-way ANOVA with Sidak’s multiple comparison tests were used, as indicated in figure legends.

### Data availability

The complete ICP0 gene sequence of the HSV-2 strain 186 has been deposited to the Gene database (https://www.ncbi.nlm.nih.gov/nuccore) and assigned the accession number MT919989. Raw high-throughput sequencing data for the PAR-CLIP experiment have been deposited to the Gene Expression Omnibus (https://www.ncbi.nlm.nih.gov/geo) and assigned the identifier GSE155402.

## Acknowledgments

The work started in Donald M. Coen laboratory at Harvard Medical School. We thank Donald M. Coen for advice and Victoria A Parsons for preparative work including some cloning and sequencing. We thank the Core Facility of Zhejiang University School of Medicine and the Biopolymers Next-Gen Sequencing Core Facility at Harvard Medical School for expertise and instrument availability. This work was supported by the National Natural Science Foundation of China (81671993 to D.P.); the Natural Science Foundation of Zhejiang Province (LR18H190001 to D.P.); the National Key R & D Program of China (2017YFC1200204 to D.P.); the National Institute of Health (P01 AI098681, for support of Donald M Coen laboratory). The funders had no role in study design, data collection and interpretation, or the decision to submit the work for publication.

## Author Contributions

D.P. conceived the study. S.C., Y.D., X.Y., B.S., and Y.L. performed molecular cloning. S.C., H.C. and D.P. performed experiments in cell culture. S.C. and D.P. generated recombinant viruses. S.C. conducted the experiments using mouse primary neurons. B.S. and Y.D. constructed knockout cell lines. S.C. and D.P. prepared the manuscript. All authors revised and approved the manuscript.

